# Detecting misfolded non-covalent lasso entanglements in protein structures, simulation trajectories, and mass spectrometry data

**DOI:** 10.64898/2026.04.15.718775

**Authors:** Ian Sitarik, Yang Jiang, Hyebin Song, Edward P. O’Brien

## Abstract

A previously overlooked class of protein entanglements, non-covalent lasso entanglements (NCLEs), has been found to play a role in widespread protein misfolding. However, understanding the influence NCLEs have on biological processes is hindered by the absence of dedicated algorithms and computational tools to detect and characterize these geometries in protein structures, molecular dynamics simulations, and in comparison to experimental data from limited proteolysis (LiP) and cross-linking (XL) mass spectrometry (MS). Here, we present EntDetect, a software tool designed to: (1) identify non-redundant NCLEs in protein structures, (2) detect misfolded states by comparing NCLE changes through pairwise comparisons of structures, (3) extract structural ensembles consistent with experimental signals from LiP-MS and XL-MS, and (4) investigate proteome-wide protein misfolding using high-throughput MS data. We demonstrate the utility of EntDetect on a simulated structural ensemble of phosphoglycerate kinase (PGK), alongside corresponding LiP- and XL-MS experimental data. Additionally, we detail the application of EntDetect to detect misfolding associated with native NCLEs on a proteome-wide MS dataset and select candidate proteins for further investigation. This protocol is intended for biophysicists, structural biologists, and molecular biologists with domain knowledge of protein structure, mass spectrometry proteomics data, and beginner experience with Python who want to interpret their experimental observations and computer simulations results through the presence and potential misfolding of NCLE topologies. EntDetect is open-source and freely available (https://github.com/obrien-lab-psu/EntDetect). NCLEweb is also available which is a webserver that identifies NCLEs within a given user-uploaded structure (https://www.ncleweb.org/).

## Introduction

Non-covalent Lasso Entanglements^1–4^ (NCLEs) are formed when pairs of residues within a protein come into contact, interact through non-covalent interactions, and form a backbone loop that is threaded by either or both termini (Fig. 1). NCLEs have recently been suggested to cause wide-spread protein misfolding^3–5^ through either the failure to form a native NCLE (Fig. 1c,d) or gains of NCLEs (Fig. 1e,f) that are not present in the reference structure (most often the native state). Either of these changes in entanglement status can result in off-pathway, kinetically trapped intermediates that in many cases must unfold for the protein to reach its native state. NCLEs are common in native structures – they are present in the majority of globular proteins^6^, and are structurally distinct from knots^7–9^, slipknots^10^, and covalent lassos^11–14^. NCLE misfolding can explain a range of observations from the past several decades, including how synonymous mutations can alter long timescale enzyme structure and function^4^, why some misfolded proteins can bypass the refolding action of chaperones^5,15^, and, it has been argued, this class of misfolding might even contribute to disease and aging processes. Given this, research in this area would be accelerated by having analysis tools that can detect and characterize these states. Here, we present a set of protocols for investigating the role of NCLEs in protein misfolding, ranging from analyses of individual protein structures and simulations supported by experimental data to proteome-wide studies where statistical power may be more limited (Fig. 2).

**Figure 1:**
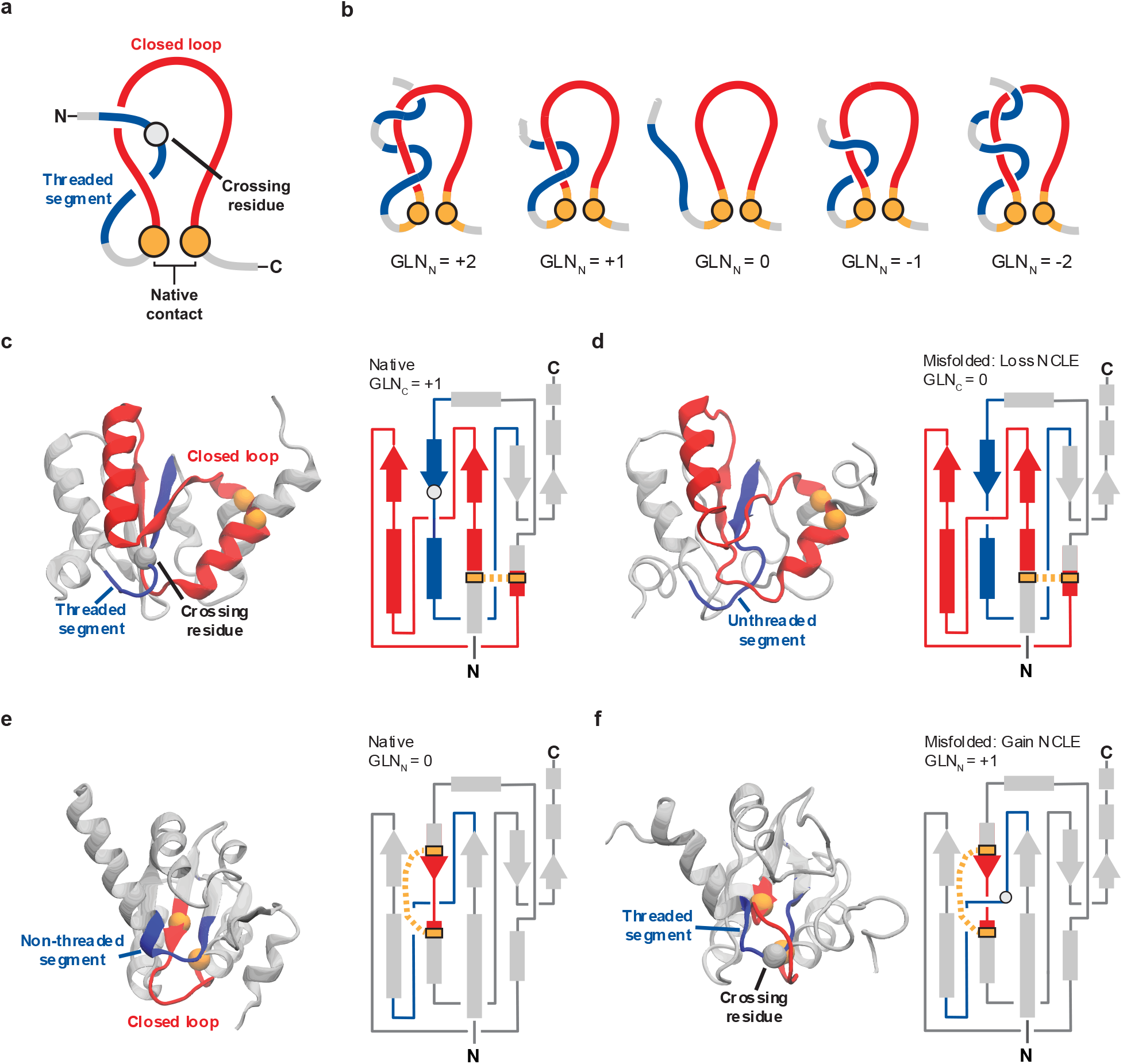
Quantification of entanglements in protein structures. **a**, The three components of a NCLE are (*i*) a loop (red) that is (*ii*) closed by a non-covalent contact (gold), defined by residues with any heavy atoms within 4.5 Å of each other, which is (*iii*) threaded by a segment outside the loop region (blue). The crossing residue is the residue on the threading segment that pierces the plane of the loop. **b**, Examples of entanglements with 1 or 2 crossings in the N-terminus and differences in the chirality of the rounded Gauss linking number (GLN) **c**, The crystal structure and 2D schematic of the large ribosomal subunit protein uL10 (P0A7J3, PDB 6XZ7, chain H) with a native NCLE consisting of a loop (red) closed by residues V12 and L59 (gold spheres – C_*α*_ atom in space fill) that is threaded by a C-terminal segment (blue) with a crossing at S85. **d**, Same as in **c** but for a highly native misfolded structure (*Q* = 0.991) where the native NCLE was lost. **e**, The crystal structure and 2D schematic of the large ribosomal subunit protein uL10 (P0A7J3, PDB 6XZ7, chain H) with a loop (red) closed by residues Y83 and L92 that is not threaded and has a 0 linking number. **f**, Same as in **e**, but showing the same highly native misfolded structure as in **d**, but which gained a NCLE by threading the loop by the N terminus with a crossing at S23.

**Figure 2:**
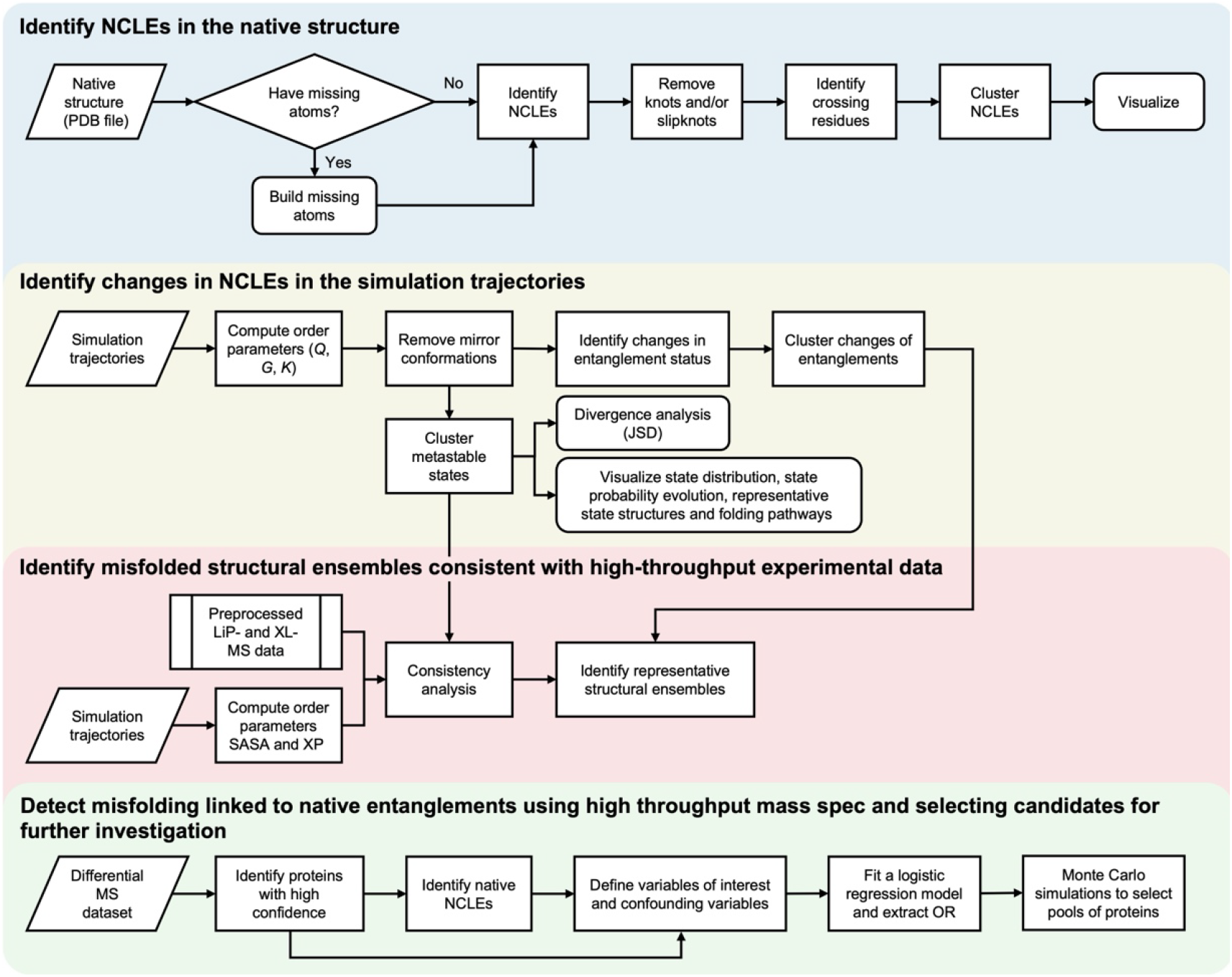
A schematic overview of the pipeline to analyze native entanglements and their changes in simulation data and mass spectrometry. All analysis in this protocol requires at a minimum the non-covalent lasso entanglements of a reference structure or state (often referred to as the native entanglements if the “native” structure is used). From simulation data the changes in the status of NCLEs can easily be calculated and in combination with high dimensional clustering techniques a set of metastable populations of structures can be identified. These metastable populations can then be compared to experimental data probing structural changes (such as LiP- and XL-MS) to quantify the consistency between simulation and experimental. Candidates for further experimental interrogation can be selected through a Monte Carlo style selection which exploits population statistics to skirt poor individual protein statistics.

### Development of the protocol

Protein entanglements, including knots, lassos, and links, have been extensively studied^8,16^. Over the last six years, our group has focused on an overlooked subtype of lassos closed by non-covalent interactions, investigating their prevalence^6^, functional implications^3,4^, evolutionary significance^6,17^, and association with protein misfolding^3,4,18–22^. To facilitate these studies, we developed a structure-based metric *G* (Eq. B1.2) which is the fraction of native contacts that exhibit a change in entanglement status^3,4,20^, enabling clear differentiation between misfolded and native states. Additionally, we established clustering workflows to identify representative, non-redundant non-covalent lassos in native structures^6^ and track changes in entanglement status with time^22^. Building upon this foundation, we developed a comprehensive analysis protocol for individual protein structures and molecular dynamics (MD) simulation trajectories that captures heterogeneous structural ensembles, elucidates folding and misfolding pathways, assesses structural ensemble divergence between conditions, and tests for consistency and statistical associations with experimentally observed structural changes. These protocols have been successfully applied to investigate the prevalence and properties of native NCLEs across species^6,17^, their misfolding dynamics^3,4,18–20,22,23^, elucidate the impact of synonymous mutations on enzyme function^4,20^, and to explain the unusual refolding kinetics of phosphoglycerate kinase (PGK)^22^.

### Applications of the protocol

The applications of this approach can be used to detect misfolded structures containing topological gains and losses of NCLEs in simulations, support design of novel therapeutics, and identify misfolded structural ensembles that can explain much of the mass spectrometry observed structural changes.

#### Detecting entanglement misfolded states

Traditional simulation order parameters (e.g., RMSD and fraction of native contacts, *Q*)^3^ can miss or misclassify misfolded protein states that preserve near-native contact patterns while harboring non-native geometric or topological changes, leaving a key gap in our ability to accurately identify misfolding events within MD trajectories. This protocol addresses that gap by tracking changes in NCLEs, providing a sensitive, geometric signature that exposes misfolded and metastable conformations that conventional metrics are blind to during folding, unfolding, and functional transitions. This is significant because it enables the community to discover new misfolding pathways and kinetic traps^4,22^ with a topology-aware descriptor, improving mechanistic interpretation of MD data^3–5,15,21,22^.

#### Therapeutic Design for Protein Misfolding

Designing small molecules to correct a protein’s misfolding-induced loss-of-function requires mechanistic insight into which non-native states are populated and biologically relevant. Identifying these misfolding pathways, transient intermediate states, and kinetic traps are difficult to resolve experimentally, making it challenging to identify actionable targets. This protocol provides tools to theoretically assess potential targets by coupling the tracking of modulations in the proteins folding landscape and pathways in the presence of various ligands during molecular simulations^24^ with the detailed mechanistic descriptions the NCLEs and their changes provide. The approach supports more rational design of molecules that correct protein misfolding-driven dysfunction, even when direct experimental characterization of the misfolded states is not possible or limited.

#### Predicting MS-detectable structural rearrangements

Mass spectrometry–based structural proteomics experiments (e.g., LiP-MS and XL-MS) are often used to probe changes in protein structure, but it can be hard to determine whether a given MD simulation actually provides a confident mechanistic explanation for those observables. By tracking changes in native NCLEs across simulation trajectories, the protocol identifies regions of conformational variability or flexibility that may correspond to protease-accessible or cross linkable sites^5,22,23^. These dynamic entanglement changes serve as mechanistically meaningful indicators of local or global structural rearrangements, improving interpretation of proteomics experiments and supporting the generation of testable hypotheses about proteomic observables.

### Comparison with other protocols and methods

We compare our approach to established structural analysis methods used to study protein topology, folding/misfolding, and experimental comparison.

#### Knots and covalent lasso entanglements versus NCLE

Our method is focused on NCLEs instead of the long-studied presence of knots^25–28^ and covalent lasso entanglements^11–13,29^ (CLE) because NCLEs are far more common. Knots and CLEs combined are represented in ∼20%^6,8^ of the known structures of the PDB while NCLEs are represented in 65.8%, 62.3%, and 54.0% of the known structures of the common model organisms proteomes, *E. coli, S. cerevisiae*, and *H. sapiens* respecfully^6^. These numbers approach 70% when high-quality Alpha-Fold structures are utilized covering a larger proportion of the proteome. With this high prevalence, NCLEs represent an easily understood structural feature that can be probed for influences on function and misfolding without the extra constraints of having to form disulfide bonds as in covalent lassos.

#### Clustering of simulation structures into metastable states

This method clusters simulation structures as a function of the fraction of native contacts (*Q*) present and the fraction of native contacts that exhibit a change in NCLE linking number (*G*, Eq. B1.2) in order to generate the set of microstates that are then clustered into metastable states using Markov state modeling^30^ (MSM). Other methods will generally either identify esoteric order parameters through manual selection or data driven identification (e.g., principal component analysis, PCA) that are often uninterpretable and system-specific^31^. Conversely there are a variety of general order parameter such as RMSD that is interpretable but has low resolution and high structural degeneracy^32^. This method uses the easy to interpret order parameters *Q* and *G* allowing for robust identification of near-native like misfold structures that would be hidden in simpler order parameters like RMSD.

#### Comparison to experiment versus integration

Rather than incorporating experimental data directly into the force field or sampling protocol through biased schemes such as MELD (Modeling Employing Limited Data)^33^ or experiment-directed simulation (EDS) approaches that add restraint potentials to enforce agreement with ensemble-averaged observables^34,35^, this method leaves the simulations unbiased and instead performs a rigorous, post hoc comparison between simulated ensembles and experimental signals. In doing so, it emphasizes statistically well-founded tests of agreement.

### Limitations

While our method provides broad utility for characterizing entanglement misfolded protein structures and comparing them against experimental data, several important limitations should be kept in mind when interpreting the results. In particular, the methods applicability is limited to the most common structural representations, and its performance depends on the statistical power and quality of the experimental datasets used for comparison, the finite resolution of the *Q*/*G*-based clustering into metastable states, and occasional false-positive NCLE detections within a given protein conformation. Below, we outline these constraints and suggest practical strategies for mitigating their impact in typical applications.

#### Model resolution

Our protocol is designed to analyze protein structure representations that report coordinates of the alpha carbon atom of each residue. However, when applied to the rare situation of coarse-grained models that exclude alpha carbons (e.g., side-chain only models^36^), the protocol is unable to identify entanglements or perform downstream analyses due to a violation of core assumptions of the method. To address this limitation, users can attempt to back-map the high-level coarse-grained model to predict alpha carbon positions using the tools such as cg2all^37^.

#### False entanglements

There is a rare occurrence of false positives in the identification of NCLEs, i.e., NCLEs are identified that are not actually present in the protein conformation. We recommend addressing this limitation by first, minimizing false positives through optimization of variable parameters in the Gauss linking integration^1–3,6^ that is the core of the protocol relative to a hand curated ground truth dataset (we recommend at least 20 proteins). Specifically, the rounding threshold used to approximate Gauss Linking Numbers^5,6^ (GLN), buffers between the loop and threading segments, and the distance a crossing can be from the loop closing or another crossing. Then, by explicit conformation of entanglement thread piercings of the loop using the Topoly^38^ python package. We finally recommend that users manually inspect random samples of their detected entanglements or changes in entanglement by visualizing the structures to ensure the method is providing accurate identification.

#### Statistical power of experimental data

The quality of the experimental data and how it is processed will heavily influence the resulting comparison with simulated misfolded protein ensembles. Here we handle cases where there are high quality experimental data measuring conformational changes for a single protein as well as proteome-wide experimental data where the statistical power is much less on the per protein level. Thus, this method is not designed to analyze experimental datasets of individual proteins with low statistical power. To address this limitation, users should obtain higher quality experimental data. We often find that 3 technical replicates are not sufficient even for single proteins and recommend between 5 to 7 technical replicates across a minimum of 4 biological replicates when performing LiP-^39^ or XL-MS.

#### Clustering of simulation structures into metastable states

The order parameters *Q* and *G* were chosen to balance interpretability and resolution and as with most projections from a high-to low-dimensional space degeneracy occurs. There indeed may be metastable states which contain structures with distinct changes in entanglement but very similar *Q* and *G* values. Improving the resolution while maintaining physical interpretability of the resulting clusters is an ongoing endeavor, and will likely be question, project, and even system specific. To address this limitation a higher resolution approach can be taken in which the user can cluster the microstates along both *Q* and the pre-clustered changes in entanglement calculated in this method (see Protocol Step 11).

### Protocol overview

#### Characterizing Protein Entanglements in a Single Structure

Any structure derived from experimental or theoretical methods can be examined for the presence of a NCLE (Fig. 1), as long as the structure has alpha carbon atomic positions reported. The native contact closing a loop in the structure can be defined in two common ways as either those residues with less than or equal to 8 Å between their alpha carbons, or when higher resolution structures are available, as those residues with less than or equal to 4.5 Å between any heavy (non-hydrogen) atoms. To determine if the loop closed by residues *i* and *j* is threaded (i.e., entangled) with a portion of the N- or C-terminus spanning residues *m* and *k*, we use a discrete version of Gaussian linking integration (Box 1) to determine the partial Gaussian linking value (*g*(*i, j, m, k*), Eq. B1.1). Since we examine one closed curve (loop) and an open curve (thread) the accuracy of *g*(*i, j, m, k*) in determining the linkage is diminished and can produce some small number of false positive NCLEs. To mitigate these, we ignore the first and last 5 residues of the protein primary structure in the analysis and the 4 residues proceeding the first residue in the loop closing contact (often indexed as *i*) and after the last residue closing the loop (often indexed as *j*)^3,4^. Second, rigorous comparisons to a human curated dataset of entanglements determined that a threshold of |*g*(*i, j, m, k*)| ≥ 0.6 for determining the presence of an entanglement balances the rate of false positive and false negative entanglements most effectively^6^. Furthermore when we examine the probability of finding a true entanglement with an orthogonal more computationally expensive but accurate method (Topoly^38^), we still find 0.6 is a reasonable cut off to mitigate false positives and negatives (Supplementary Figure 1). Therefore, when determining the GLN (Fig. 1b, Box 1) we apply a modified rounding rule in which values are rounded up or down depending if the remainder absolute value is less than or greater than 0.6, rather than the conventional 0.5 threshold. It is worth noting that many PDB structures have missing residues and they can be rebuilt using MODELLER^40^ or CHARMM^41^, or entanglements involving the missing residues need to be removed from the analysis (Box 1).

Once a loop is identified as being entangled, finding the locations along the thread where the loop is pierced can provide biophysical insight into a proteins’ folding and function^4,11,20^. One fast but less accurate approach to identify these crossing residues is to examine the behavior of |*g*(*i, j, m, k*)| as a function of a sliding window of 15 residues along the terminal tail of interest^20^. This function will reach a maximum at the neighborhood where the thread pierces the plane of the loop. A more accurate, yet slower, approach is based on tessellating the surface of the loop using a series of triangular planes that can efficiently identify the crossing residue by examining which bond vectors cross them and is available in the software package Topoly^38^. It is in general recommended to use the tessellation approach, when possible, but it may become a computational bottleneck when thousands of structures need to be analyzed.

A single protein structure may have many degenerate entanglements that have loop closing contacts and crossing residues close along the primary structure and identical topological linkage numbers. We therefore developed an algorithm to cluster loops with crossing residues that are spatially close, and which share the same crossing chirality (Box 2). The output of this clustering is a representative entanglement for each cluster consisting of the minimal closing loop and the crossing residues located on the N- or C-terminal thread(s). These representative entanglements are best visualized in VMD^42^ or PyMol^43^, and we recommend a community standard (see Fig. 1) in which the cartoon representation of the loop and thread are colored, respectively, red and blue. The alpha carbons of the loop closing contacts and crossing residue(s) should be in space filled representation and colored orange and white, respectively (e.g., see Fig. 1).

#### Detecting Non-Native Entanglements in Simulation Trajectories

Changes of entanglement status in simulation trajectories, corresponding to either gains or losses of NCLEs (Fig. 1d,f), are characterized by the order parameter *G* (Eq. B1.2, Box 1) which captures the fraction of native contacts with changes in linkage between a given simulation structure and some reference, often chosen as the protein’s native structure^4^. A larger value of *G* indicates more loops had a change in entanglement status and when *G* = 0 then there is either no change in topology relative to the reference structure or no native contacts have been formed. Mirror artifacts^44–46^ (or images) in the simulation trajectories where the packing chirality of secondary and tertiary structure is reversed can inflate *G*, and they must be excluded from the entanglement analysis, especially in models with highly isotropic and symmetric interaction potentials and structures. Such mirror artifacts, while uncommon in coarse-grained models, are very rare in all-atom representations. An order parameter *K* (Eq. S2 in Ref. 12) can be used for this purpose. It quantifies the fraction of tertiary structure chirality that is identical to those in the native structure^47^, which is 1 when correct chirality is maintained and 0 when a mirror image is present^22^. Simulation trajectories with an average fraction of native contacts (*Q*) > 0.2 and *K* < 0.6 are flagged as artifacts and should be visually verified and removed from downstream analyses.

Structural distributions across the simulation trajectories can then be visualized by examining a probability surface defined by −*ln*(*P*(*Q, G*)), where *P*(*Q, G*) is the probability density of simulation conformations characterized by *G* and *Q*, with a number of microstates (e.g., 400) clustered using k-means and further grouped into metastable states via Markov state modeling^30^ (MSM) (Fig. 3a). The folding pathways (Fig. 3g and h) can then be determined by tracking discrete metastable-state transitions, allowing for the identification of transitions into and out of entangled misfolded states, as well as state distributions and transition rates^4^. It is often useful to have a single representative structure of each metastable state which can be randomly sampled from all microstates according to the probability distribution of the microstates within the given metastable state^4^. These analyses have been applied to identify intermediate states that can be potential drug targets to prevent entanglement misfolding^24^.

**Figure 3:**
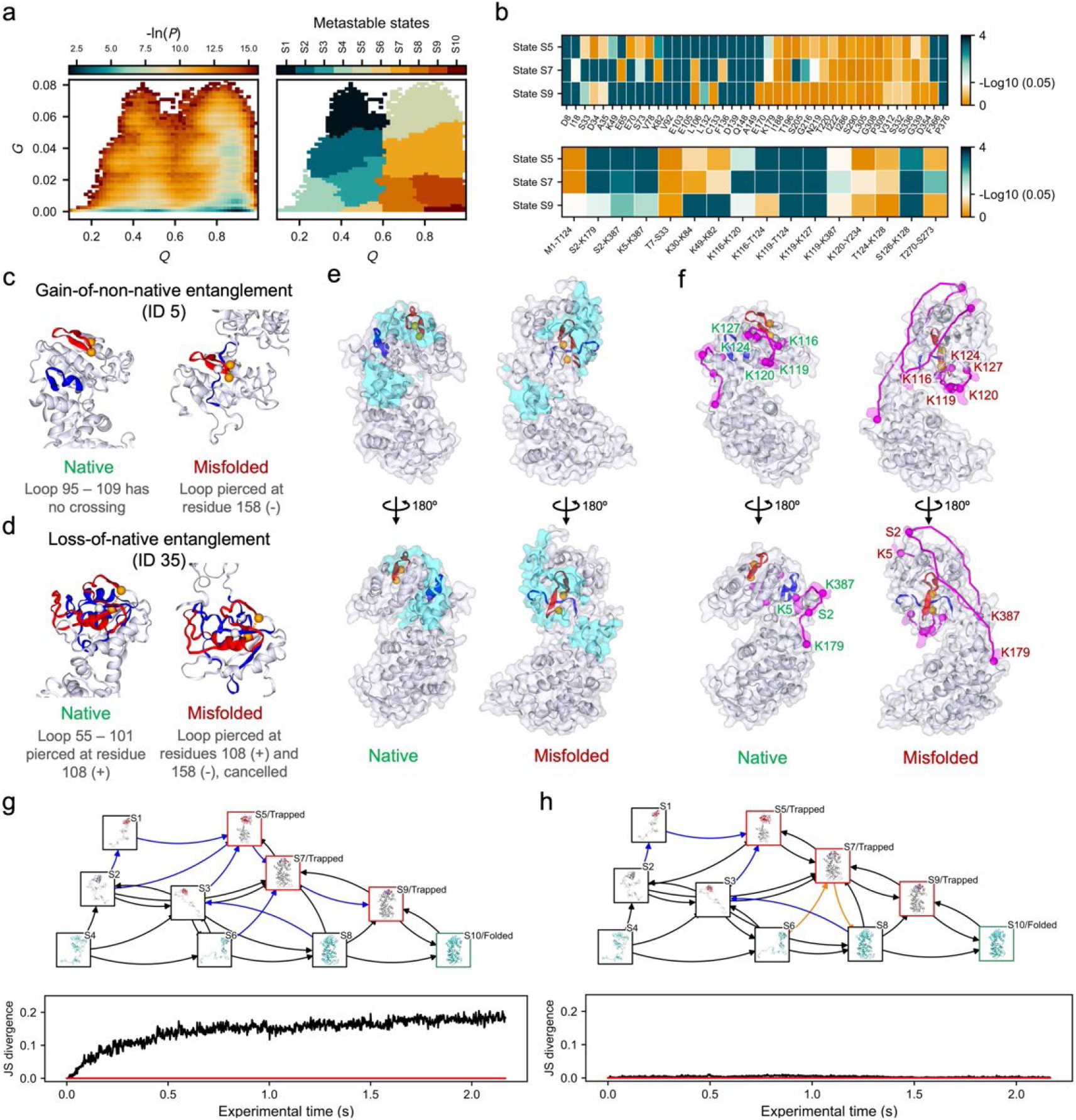
Simulation trajectory analysis and comparison of misfolded structural ensemble with LiP-MS and XL-MS data. **a**, The results of building a MSM across the full quenched trajectories. The -ln(P) surface (left) and the identified metastable states (right). **b**, −log_10_(adjusted P values) for the test of consistency between the structural changes observed in the near-native misfolded states S5, S7, and S9 (regarding the native state S10) against those suggested by LiP-(top) and XL-MS (bottom) data. **c** and **d**, the representative changes in entanglement found in state S5. **e**, consistent LiP-MS signals highlighted on the native structure (left) and misfolded structure (right). Entanglement changes ID 5 is highlighted. PK cut-sites are represented as cyan balls and residues (±5) around the cut-sites are transparent cyan surfaces. **f**, consistent XL-MS signals highlighted on the native structure (left) and misfolded structure (right). Entanglement change ID 5 is highlighted. Residue pairs that identified in XL-MS signals are shown as magenta balls with transparent magenta surface and SASDs identified by Jwalk as magenta curves. **g**, folding pathways (top) and JSD vs. time (bottom) of ecPGK under two artificial conditions, where the trajectories were divided into two groups, one with 80% trajectories converging to the native state and the other with 20%. **h**, folding pathways (top) and JSD vs. time (bottom) of ecPGK under another two artificial conditions where the trajectories were randomly split into two groups. The folding pathways are presented as a directed graph where the nodes represent metastable states, with representative structures shown in the boxes. The edges represent the transitions in the pathways. The nodes corresponding to the native state and the kinetically trapped states are framed in green and red, respectively. The transitions (arrows) that are observed in only the condition A and condition B are marked in orange and blue, respectively. The red line in the JSD plot represents JSD = 0, i.e., totally converged between two conditions.

The observed changes in entanglement will have some degree of degeneracy and it is helpful to identify representative entanglement changes across simulation structures. A simple clustering and categorization algorithm^22^ is applied by first categorizing the NCLE changes based on its type (e.g. the nomenclature ‘L+C#’ means a gain of linking number without a change in crossing chirality, Box 2). Clustering analyses are then performed based on crossing residues, loop location, and crossing contamination (Box 2). The permuCLUSTER algorithm^48^ was used with 100 permutations to mitigate input order bias, and the result is a set of unique change in entanglement status with different types and locations on the protein (examples are shown in Fig. 3c and d).

Since some of these misfolded states are predicted to be long lived, and kinetically trapped, it can be of interest to explore the impact of two different initial conditions on the resulting misfolded ensemble and the associated hysteresis that can arise^4^. In this case, comparing the time evolution of the two resulting ensembles can be carried out at a given time point using the Jensen-Shannon divergence (JSD) metric^49^:

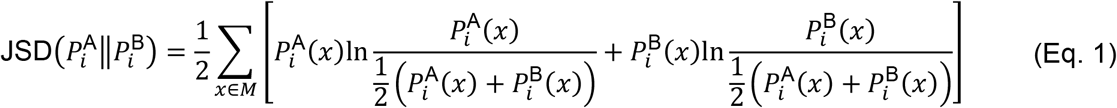

where 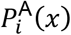 and 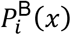 are the probability of the metastable state *x* at time point *i* under condition A and B, respectively. This approach was recently used to characterize the impact of changes in translation speed on these structural ensembles^4^, where A = fast translation and B = slow translation, and provides a useful tool to understand the influences of different initial conditions on the formation of non-native NCLE (examples generated by this protocol can be found in Fig. 3g and h).

#### Identifying Entangled Structural Ensembles Consistent with High-Throughput Experimental Data

Two common high-throughput proteomics techniques probing protein structure changes are cross-linking^50,51^ (XL) and limited proteolysis^52,53^ (LiP) mass spectrometry (MS). Both methods can detect changes in protein structure by comparing peptide abundances under two different conditions (i.e. treated and untreated samples). LiP-MS relies on comparing changes in protease accessibility as a proxy for structural change and XL-MS relies on comparing changes in the frequency in which two cross-linkable residues were within a certain distance threshold of each other. The structural data generated can constrain the potential structural ensembles that exist and comparing the consistency between this experimental data derived from samples containing purified protein and ensembles of protein structures generated through simulation or other means can provide useful biophysical insights.

To assess the consistency of a metastable state in the ensemble of simulation structures with LiP-MS experimental data, the location of structural changes observed in the experimental data must first be identified. We find the proteinase-K (PK) cut-sites on half-tryptic peptides that have a statistically significant change in their abundance after refolding and false discovery rate (FDR) correction (Benjamini Hochberg, *α* = 0.05)^54^ hold the most physically interpretable meaning. To connect to simulated ensembles we assume that there must be a change in solvent accessible surface area between the simulated misfolded ensemble and the native state for there to be any change in protease digestion patterns. We use the list of experimentally observed cutsites and for each calculate the difference in solvent accessibility, including +/-5 residues around it, between the metastable state and the native ensemble. We test if this difference is significant using a two-tailed permutation test and an FDR correction. If a cutsite exhibits a significant difference in solvent accessibility we consider it consistent with the experimental observation. The simulated metastable states exhibiting the highest number of consistent cutsites are identified as the most consistent structural ensemble with the experimental protease digestion pattern (Fig. 3b top).

In XL-MS experiments the location of structural changes are identified similarly by the pair of cross-linked residues that have a statistically significant change in their abundance even after FDR correction (Benjamini Hochberg, *α* = 0.05). We then test the folding simulations for similar changes in crosslinking propensity (XP) between misfolded and native states. XP was estimated using a modified MNXL scoring function^22,55^, which accounts for solvent-accessible surface distance^55^ (SASD) and was adjusted for non-Lys residues and a longer crosslinker molecule disuccinimidyl dibutyric urea (DSBU). As above, for the list of observed cross-linking residues, if the difference in the simulated XP between the mestastable state and the native ensemble is significant we consider it consistent with experiment. The metastable states with the highest number of consistent cross-linked pairs are the most consistent structural ensemble with the XL-MS data (Fig. 3b bottom).

The statistical test described above evaluates the consistency at the ensemble-averaged level. To select representative misfolded structures from each metastable state that exhibit structural changes consistent with MS observations – for either visualization or downstream, higher-resolution structural analyses – simulation structures are first grouped according to the clusters of entanglement changes (as defined by the cluster IDs). They are then further grouped by their list of cutsites and cross-liked pairs whose SASAs and XPs show statistically significant changes compared with the native ensemble (values falling outside the 95% CIs of the native ensemble)^22^. Within each resulting group, the structure with the highest microstate probability is selected as the representative, forming the final representative misfolded structural ensemble (Fig. 3e and f).

#### Detecting misfolding involving native entanglements using high-throughput mass spectrometry and selecting candidates for further investigation

When examining multi-protein samples with LiP-MS, such as in the case of whole or sub proteome samples, there is often a significant reduction in statistical power to detect conformational changes in individual proteins relative to single-protein samples. This can make it difficult to detect signatures of entanglement misfolding but there are several strategies that can be used to address challenge.

First, some statistical power can be regained by the application of filters to remove poorly sampled proteins. For proteome-wide experimental datasets, only the proteins with at least 50% of their canonical sequence detected in the untreated group should be included for further analyses. This limits false negatives in downstream analyses and proteins that are inherently difficult to observe via mass spectrometry. Second, it is also useful to control differences in the natural abundance of proteins in the sample. The sum of the peptide abundance (SPA, Box 4, Eq. B4.1) is a decent proxy for the natural protein abundance^56^ and can be used to threshold the dataset or test the robustness of results against.

Further statistical power can be gained by aggregating data across subsets of proteins (i.e., population level behaviors). Applying these three approaches, it is possible to examine whether underlying associations exist between LiP-MS observed changes in conformation and the presence of native NCLEs, as well as their location along the primary structure. To obtain such associations, Logistic regression^57^ is commonly used as it can simultaneously control for multiple confounding factors and it computes the log-odds of a protein misfolding as a function of the presence of a native entanglement and protein (Box 4, Eq. B4.2). The magnitude and direction of the association of misfolding and the presence of native entanglements is measured by the coefficients in the logistic regression fit to the data, which can be transformed into an odds ratio (Box 4). At the residue level, we can measure the misfolding bias in a particular region along a protein’s primary structure by modeling the log-odds of a specific residue showing a significant change in abundance as a function of the location of the residue in the protein structure (in our case whether it was in a natively entangled region or not) and the confounding factors of amino acid composition and solvent accessibility (Box 4, Eq. B4.3).

Due to the lack of statistical power at the per protein level in these large high throughput experiments we have found that we cannot rank order the proteins based on their likelihood to misfold involving native entanglements with any statistical certainty. Therefore, a Monte Carlo based algorithm was created to select subsets of proteins that collectively exhibit the most extreme misfolding biases with the aim of identifying potential candidates for follow up experiments or simulation studies. We initially split our set of observed proteins into *n* groups (*n* = 4, we do not recommend going below *n* = 3) and for each group we measure the strength of the misfolding bias from a logistic regression and an objective function is used to calculate a characteristic “energy” of the system (Fig. 4a, Box 4, Eq. B4.4). We then randomly swap the proteins between the ordered pairs of groups, recalculate the “energy” and apply the Metropolis criteria^58^ to determine whether we accept or reject the swap constituting a single Monte Carlo step (Fig. 4b). After extensive iterations (200,000 steps or more) we rank order the groups based on the magnitude of their misfolding bias and use the highest rank group to select candidates for further experiments or simulations.

**Figure 4:**
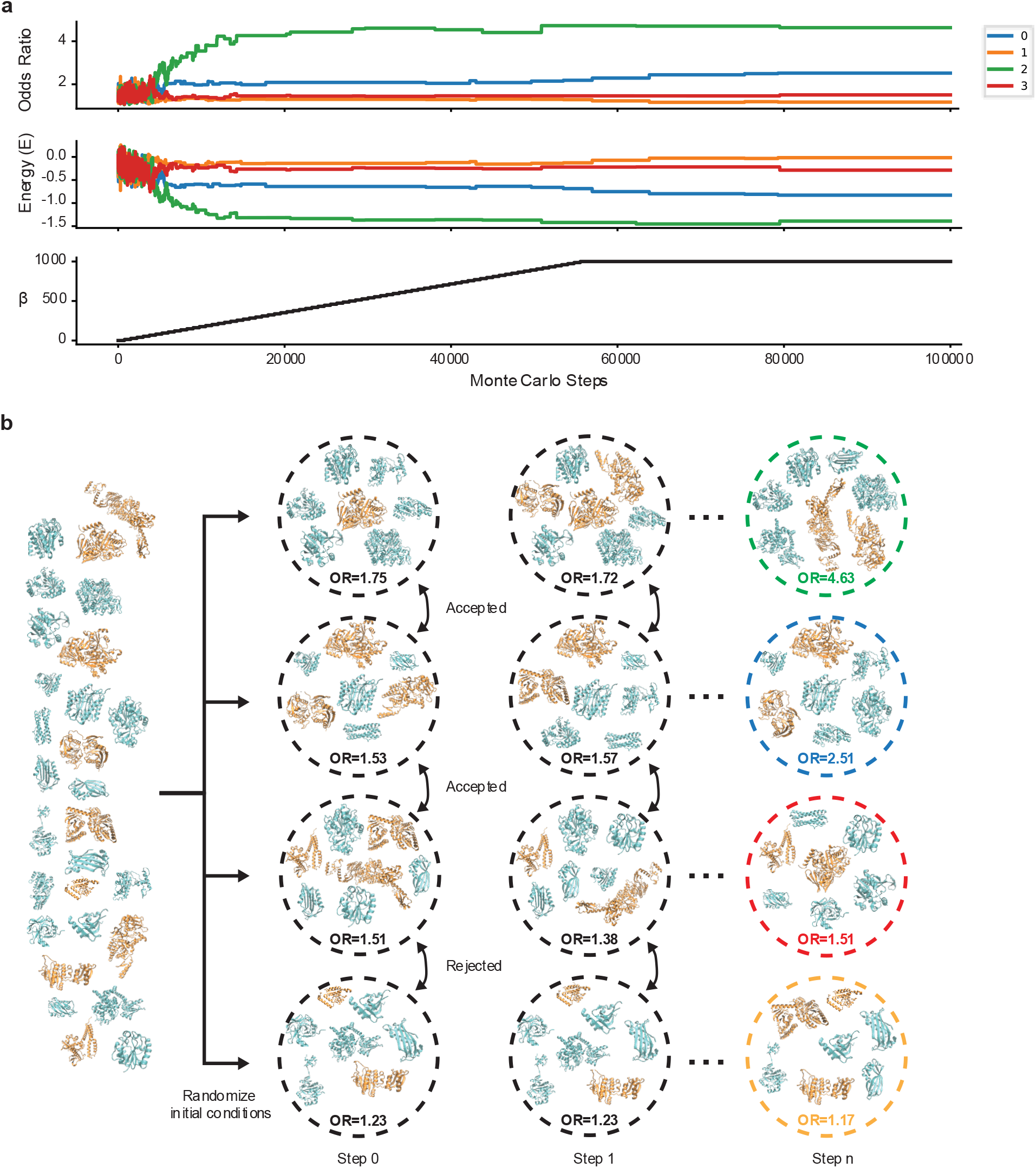
Monte Carlo style selection of candidates for simulation or further experiments. **a**, (top) the odds ratio (OR) of the association between the structural changes observed in mass spec data (in this case LiP-MS) and the region of the protein it was found (entangled region or non-entangled region). (middle) the objective function used in the Metropolis criteria at each Monte Carlo step. (bottom) a hyper parameter that controls the information “temperature”. **b**, a schematic of the Monte Carlo scheme showing how an initial collection of proteins with native entanglements is randomly placed into one of four groups and then over the course of the Monte Carlo swaps results in groups with very different populations levels of association.

## Materials

### Hardware

- The minimal compute requirements for this protocol are as follows: a Linux workstation or cluster equipped with at least 1 central processing unit (CPU) processors, 10 GB random access memory (RAM), and enough disk space for all input data and outputs.
- Performance will vary on the basis of system configuration. For compute-expensive steps of the protocol, we provide approximate timings using a system equipped with 8 CPU processors and 100 GB RAM.

### Software

- EntDetect: Download, installation guide, and tutorials are available at (https://github.com/obrien-lab-psu/EntDetect).
- Python 3.X, with numpy, scipy, matplotlib, biopython, pandas, MDAnalysis, numba, topoly, geom_median, seaborn, scikit-learn, network, pyyaml, tqdm installed. We recommend using conda to install and manage python environment. The environment.yml file in the EntDetect GitHub repo can be used install all required packages.
- VMD: Download find installation guide at https://www.ks.uiuc.edu/Development/Download/download.cgi?PackageName=VMD.

### Datasets

- Simulation trajectories: We provide the example coarse-grained simulation trajectories obtained from our previous work on PGK from E. coli (ecPGK)^22^ for users to test the procedure described in this protocol (see Procedure). They can be downloaded from <u>10.5281/zenodo.18232475</u> Note well that the protocol is valid on any other simulation trajectories regardless of the model resolution, including all-atom resolution^21^.
- Pre-processed LiP- and XL-MS data for ecPGK: Download from (https://github.com/obrien-lab-psu/EntDetect).
- Miscellaneous data required for running the procedure: For example, the PDB file of the crystal structure of ecPGK, and the secondary structure elements of ecPGK, etc. Download from <u>10.5281/zenodo.18232475</u>.
- Proteome wide LiP-MS data: Download from https://doi.org/10.1038/s41467-025-66236-3

### Protocol

This protocol assumes that you are working in a Linux environment, have installed the necessary software detailed in the materials section, and created a mini-conda environment with the EntDetect package installed. Detailed tutorials and documentation of all code in this protocol can be found on the EntDetect GitHub.

#### Identify NCLEs in the native structure and their associated features

Timing (<1min to 10min) depending on protein size

1. Enter the following at the terminal to enter the base directory and activate your conda environment with EntDetect installed. ~~~
cd /path/to/base/directory
conda activate EntDetect_env
~~~
2. Process the Protein Data Bank or AlphaFold structure file to rebuild missing residues and atoms using Modeller^59^ or Charmm^60^ and ensure there are no duplicate residues.
3. Open an interactive Python session (or add the following steps to a python script or function) and execute the following commands to identify NCLEs in the structure (Box 1).
4. Import, Initialize, and use the GaussianEntanglement and ClusterNativeEntanglements classes from EntDetect to calculate the set of NCLEs for the reference structure (Box 1). ~~~
from EntDetect.gaussian_entanglement import GaussianEntanglement
## Define some input paths and parameters
pdb = “./1zmr_model_clean.pdb”
native_outdir = “./nativeNCLE/Native_GE”
ID = “1zmr”
## Initialize the GaussianEntanglement class
ge = GaussianEntanglement(g_threshold=0.6, density=0.0, Calpha=False, CG=False)
clustering = ClusterNativeEntanglements(organism=“Ecoli”)
## calculate the entanglements
ge.calculate_native_entanglements(clean_pdb=pdb, outdir=native_outdir, ID=ID)
~~~ Outputs a csv file containing the location of the NCLE.
5. (Optional) Remove slipknots and low-quality entanglements in AlphaFold structures. (Box 1) ~~~
## Define some input paths and parameters
native_HQ_outdir = “./nativeNCLE/Native_HQ_GE”
NCLE_file = “./nativeNCLE/Native_GE/1zmr_model_clean_ca_GE.txt”
## filter the NCLE
ge.select_high_quality_entanglements(NCLE_file, pdb, outdir=native_HQ_outdir, ID=ID,
model=“EXP”)
~~~ Outputs a csv file containing the filtered NCLEs. *Note* that if the protein structure contains portions do not present in the canonical protein sequence (i.e. chimera proteins, inserted regions, ect.) a mapping file can be supplied to select for only those entanglements that do not contain any non-canonical regions.
6. Cluster NCLEs into non-redundant representative entanglements ~~~
from EntDetect.clustering import ClusterNativeEntanglements
## Define some input paths and parameters
native_clustered_HQ_outdir = “./nativeNCLE/Native_clustered_HQ_GE”
NCLE_file = “./nativeNCLE/Native_HQ_GE/1zmr.csv”
Outfile = “1zmr.csv”
## cluster the native NCLEs and get representative NCLEs
clustering.Cluster_NativeEntanglements(NCLE_file, outdir=native_clustered_HQ_outdir,
outfile=outfile)
~~~ Outputs a csv file with the representative entanglements and their degenerate loops.
7. Import, initialize, and use the FeatureGen class from EntDetect and calculate features of the representative entanglements^5^ such as the number of super coiling events of the entangled terminus, the number of contacts the loop makes with the rest of the protein, and others which capture its complexity. ~~~
from EntDetect.entanglement_features import FeatureGen
## Define some input paths and parameters
pdb = “./1zmr_model_clean.pdb”
native_GQ_feature_outdir = “./nativeNCLE/Native_clustered_HQ_GE_features”
cluster_file = “./nativeNCLE/Native_clustered_HQ_GE/1zmr.csv”
## Initialize the feature generation class and get the features for each unique NCLE
FGen = FeatureGen(pdb, outdir=native_GQ_feature_outdir, cluster_file=cluster_file)
EntFeatures = FGen.get_uent_features(gene=‘P00558’, chain=‘A’, pdbid=‘1ZMR’)
~~~ Outputs a csv file with the features for each representative entanglement.
8. Visualize representative NCLEs in VMD or PyMol.

#### Identify changes in NCLEs in the simulation trajectories

Timing (hours to several days) depending on the number of trajectories and the size of the protein.

9 Compute the order parameters *Q, G*, and *K* for each frame of the CG trajectories
  a. Import, Initialize, and use the CalculateOP class from EntDetect to calculate the key order parameters. This will require the protein structure file (psf) and coordinate file (cor) of a reference state to compare each frame of the trajectory against, a file containing the secondary structure elements identified by STRIDE^61^, the dcd file of to be analyzed, and a file containing the CATH^62^ domain definitions for the protein. ~~~
from EntDetect.order_params import CalculateOP
## Define some input paths and parameters
Traj = 1
PSF = “./1zmr_model_clean_ca.psf”
DCD = “./1_prod.dcd”
ID = “1ZMR”
COR = “./1zmr_model_clean_ca.cor”
sec_elements = “./secondary_struc_defs.txt”
domain = “./domain_def.dat”
outdir = “./run_OP_on_simulation_traj_last67frames/”
start = 6600
## initialize the order parameter calculation class
CalcOP = CalculateOP(outdir=outdir, Traj=traj, ID=ID, psf=psf, cor=cor,
sec_elements=sec_elements, dcd=dcd, domain=domain, start=start)
~~~
  b. Compute order parameters. ~~~
## Calculate Fraction of native contacts (Q)
Qdata_dict = CalcOP.Q()
## Calculate the fraction of native contacts with a change of entanglement (G)
## with Topoly, using Calpha distances to define native contacts, and a
## coarse grained trajectory.
Gdata_dict = CalcOP.G(topoly=True, Calpha=True, CG=True, nproc=10)
## Calculate the mirror symmetry order parameter K
Kdata_dict = CalcOP.K()
~~~ Outputs a new directory for each order parameter and at minimum a time series of the metric in a csv file. For *G* there is a significant amount of metadata for each frame analyzed that is also stored in binary pkl files. *Pause Point*. The processing time can be days when the trajectories are large. You can down sample the trajectories to save analysis time.
10 Identify and remove artificial mirror conformations
  a. Apply cutoffs for *Q* and *K* by examining the steady state portion of the trajectory to identify potential trajectories with mirror images (⟨*Q*⟩ > 0.2 and ⟨*K*⟩ < 0.6). Note that these two cutoffs are chosen based on our experience on the simulated structures of ecPGK. Adjust them if you see fit in your study.
  b. Visually inspect these trajectory structures and discard those that are mirror images
11 Cluster changes of entanglements to remove redundancy.
  a. Import, initialize, and use the ClusterNonNativeEntanglements class from EntDetect. This will require the path to the binary files containing all the combined changes in entanglement information created in step 9b (`/step_9b_outdir/G/Combined_G`), a file containing the mapping between trajectory number and dcd file name, and the path to the directory containing the dcds. Run the multistep clustering algorithm across all trajectories (Box 2). Here we only cluster the last 67 frames in each trajectory (starting frame number = 6,600), which corresponds to 10 ns. ~~~
from EntDetect.clustering import ClusterNonNativeEntanglements
## Define some input paths and parameters
pkl_file_path = “./OP/G/Combined_GE/”
trajnum2pklfile_path = “./trajnum2file.txt”
traj_dir_prefix = “/path/to/dir/containing/dcds/”
outdir = “./nonnative_entanglement_clustering”
## initialize the change in NCLE clustering class
clustering_NNents = ClusterNonNativeEntanglements(pkl_file_path=pkl_file_path,
trajnum2pklfile_path=trajnum2pklfile_path, traj_dir_prefix=traj_dir_prefix,
outdir=outdir)
## Dod the clustering
clustering_NNents.cluster(start_frame=6600)
~~~ Outputs summary tables that capture representative entanglement changes, their structural fingerprints, cluster memberships, and probabilities, plus a compressed archive of all clustering inputs and mappings. It also outputs a distribution plot of loop/crossing residues, a text description of the clustering tree.
  b. Visualize non-redundant changes of entanglements *Pause Point*. The processing time can be days when the numbers of raw changes of entanglements are large. The memory consumption will be also large, so users must make sure the memory of their workstation is sufficient.
12 Build MSM to cluster metastable states
  a. Import, initialize, and run the MSMNonNativeEntanglementClustering class from EntDetect to build the Markov model. This will require the path to the order parameters from step 9b and the maximum number of metastable states to be assigned for the largest connective subgraph (n_large_states), with a higher number providing finer resolution of structural states. (Box 3) ~~~
from EntDetect.clustering import MSMNonNativeEntanglementClustering
## Define some input paths and parameters
outdir = “./run_MSM”
ID = “1ZMR”
OPpath = “./run_OP_on_simulation_traj_Allframes/”
n_large_states = 10
## initialize the MSM class and build the model
MSM = MSMNonNativeEntanglementClustering(outdir=outdir, ID=ID, OPpath=OPpath,
n_large_states=n_large_states)
MSM.run()
~~~ Outputs a csv file mapping each structure to the micro-state and meta-stable state identifiers resulting from the Markov model. Also plots the −*ll*(*P*(*Q, G*)) surface and meta-stable state map along the same surface. *Critical Step*: Try several values for n_large_states and visualize the results to select an optimal value. More metastable states can reduce interpretability, and we find that 15 or fewer usually strikes a good balance. Note that the final number of states may be lower than specified, as empty states are automatically discarded. We also by default use a lag time of 1 frame for the Markov state modeling without checking Markovian behavior. It is in general acceptable to do so if the purpose is a rough decomposition of microstates into broader regions for visualization, initial coarse-graining, or downstream experimental comparisons. Users must be cautious of choosing the lag time to make sure Markovian behavior if they want to interpret kinetics for the metastable states.
13 Visualize state distribution, state probability evolution, representative state structures and folding pathways
  a. State distribution as depicted using −*ln*(*P*), see Fig. 3a. This file is automatically generated by step 12.
  b. Plot state probability evolution along simulation time. ~~~
from EntDetect.statistics import MSMStats
## Define some input paths and parameters
outdir = “./MSM_StateProbabilityStats”
msm_meta_file = “MSM/1ZMR_prod_meta_set_A80%Native.csv”
meta_set_file = “MSM/1ZMR_prod_meta_set.csv”
tarj_type_col = “traj_type_A80%Native”
traj_type_list = [‘A’, ‘B’]
rm_traj_list = []
## initialize the MSM statistics class
MS = MSMStats(outdir=outdir, msm_data_file=msm_meta_file,
meta_set_file=meta_set_file, tarj_type_col=tarj_type_col, rm_traj_list=rm_traj_list,
traj_type_list=traj_type_list, tarj_type_col=tarj_type_col)
## Calculate state probability statistics
df = MS.StateProbabilityStats()
## Plot the results
MS.Plot_StateProbabilityStats(df=df)
~~~ Outputs a time series of the meta-stable state probabilities.
  c. Visualize representative metastable state structures using VMD^42^
  d. Compute and plot folding pathways through metastable states and Jensen-Shannon divergence (JSD, Eq. 1) to assess the divergence of structural ensembles obtained from simulations under two conditions. We show a toy example using the ePGK simulations that have been either randomly split into two artificial “conditions” (i.e. converged conditions) or split into two conditions with one being overrepresented in simulations that folded to the native state and the other overrepresented in simulations that end in a kinetically trapped misfolded state (i.e. diverged conditions). ~~~
from EntDetect.statistics import FoldingPathwayStats
## Define some input paths and parameters
outdir = “./Foldingpathway_A80%Native”
msm_meta_file = “MSM/1ZMR_prod_meta_set_A80%Native.csv”
meta_set_file = “MSM/1ZMR_prod_meta_set.csv”
tarj_type_col = “traj_type_A80%Native”
rm_traj_list = []
## initialize the clustering object
msm_data = pd.read_csv(msm_meta_file)
FP = FoldingPathwayStats(msm_data=msm_data, meta_set_file=meta_set_file,
tarj_type_col=tarj_type_col, outdir=outdir, traj_list=rm_traj_list)
## get the post-transitional folding pathways
folding_pathways = FP.post_trans()
## JS divergence
JS_divergence = FP.JS_divergence()
~~~ Outputs the folding pathways and their probabilities to a csv file as well as a JSD time series file.

#### Identify entangled structural ensembles consistent with high-throughput experimental data

Timing (∼5 – 24 hrs) depending on number of protein structures in ensemble and experimental signals.

14 Prepare experimental data files from processed LiP- and XL-MS data (Box 3).
14 (Coarse-grained C-alpha trajectories only) Back map the trajectory to the all-atom resolution
  a. For each frame in the final portion of each trajectory save a PDB file
  b. Back-map each CG PDB file to the all-atom resolution.
    i. Import, initiate, and use the BackMapping class from EntDetect ~~~
from EntDetect.change_resolution import BackMapping
## Define some input paths and parameters
Outdir = “./BackMapping/”
cg_pdb = “./1zmr_model_clean_ca.cor”
aa_pdb = “./1zmr_model_clean.pdb”
ID = “1ZMR”
## initiate back mapping class and back-mapped Calpha coarse grained structures
backMapper=BackMapping(outdir=‘./BackMapping/’)
backMapper.backmap(cg_pdb=cg_pdb, aa_pdb=aa_pdb, ID=ID)
~~~ Outputs fully reconstructed, energy-minimized all-atom protein structure whose backbone and sidechains are consistent with the supplied coarse-grained Cα model and native reference. Depending on how it’s run, it can also produce intermediate structures and logs from the PD2/Pulchra^63,64^ reconstruction and OpenMM minimization steps. *Pause Point*. Examine the backmapped structures for quality and then collate them into a single dcd for each trajectory.
16 Calculate the solvent accessible surface area (SASA), J-walk distances, and cross-linking probability (XP, Eq. B3.1) using the CalculateOP class from EntDetect. This only requires the all-atom dcd. ~~~
from EntDetect.order_params import CalculateOP
## Define some input paths and parameters
Traj = 1
PSF = “./1zmr_model_clean.pdb”
DCD = “./1_prod_aa.dcd”
ID = “1ZMR”
outdir = “./run_OP_on_simulation_traj_last67frames/”
start = 6600
## initiate CalculateOP class
CalcOP = CalculateOP(outdir=outdir, Traj=traj, ID=ID, psf=psf, cor=cor,
sec_elements=sec_elements, dcd=dcd, domain=domain, start=start)
## calculate the solvent accessible surface aread
CalcOP.SASA()
## calculate J-walk distance
CalcOP.runJwalk(‘/path/to/backmapped/pdb’)
## calculate XL probability
XPdata_dict = CalcOP.XP(pdb=‘/path/to/AA/ref/PDBfile’)
~~~ Outputs metric time series required for consistency test as a compressed binary file. *Pause Point*. The processing time can be days when the trajectories are large. We recommend only analyzing the last 50 ns and down sample the trajectory every 20 frames, which removes the autocorrelation in this example and represents the long-lived states’ ensemble of structures. Note that the 20-frame down sampling rate is chosen for the ecPGK trajectories; users should adjust it accordingly for the simulation trajectories they are analyzing to effectively remove autocorrelation.
17 Analyze consistency between simulation structural ensembles and the experimental data
  a. Import, initiate, and use the MassSpec class from EntDetect for the consistency analysis and representative structure selection. This will require the metastable state mapping file from the MSM, the accompanying microstate probability file, the processed LiP- and XL-MS data files, file containing the SASA and XP values, the non-native entanglement clusters, and finally determining which of the metastable states resulting from the MSM is the “native state” (<monsopace>native_state_idx</monsopace>) and which are the states you want to test for consistency (<monsopace>state_idx_list</monsopace>). ~~~
from EntDetect.compare_sim2exp import MassSpec
## Define some input paths and parameters
outdir = “./MassSpec_ConsistencyTest/”
msm_data_file = “./MSM/1ZMR_prod_MSMmapping.csv”
meta_dist_file = “./MSM/1ZMR_prod_meta_dist.npy”
LiPMS_exp_file = “./ecPGK_significant_LiPMS_peptide_R1_merged.xlsx”
sasa_data_file = “./run_OP_on_simulation_traj_last67frames/SASA/SASA.npy”
XLMS_exp_file = “./ecPGK_significant_XLMS_peptide_R1_merged.xlsx”
dist_data_file = “./run_OP_on_simulation_traj_last67frames/Jwalk/Jwalk.npy”
cluster_data_file =
“./nonnative_entanglement_clustering/cluster_data_topoly_linking_number.npz”
OPpath = “./run_OP_on_simulation_traj_last67frames/”
AAdcd_dir = “/path/to/backmapped/dcds/”
native_AA_pdb = “./1zmr_model_clean.pdb”
state_idx_list = [4, 6, 8]
prot_len = 387
last_num_frames = 335
rm_traj_list = []
native_state_idx = 9
start = 6600
## initializing the MassSpec class
MS = MassSpec(msm_data_file=msm_data_file, meta_dist_file=meta_dist_file,
LiPMS_exp_file=LiPMS_exp_file, sasa_data_file=sasa_data_file,
XLMS_exp_file=XLMS_exp_file, dist_data_file=dist_data_file,
cluster_data_file=cluster_data_file, OPpath=OPpath, AAdcd_dir=AAdcd_dir,
native_AA_pdb=native_AA_pdb, state_idx_list=state_idx_list, prot_len=prot_len,
last_num_frames=last_num_frames, rm_traj_list=rm_traj_list,
native_state_idx=native_state_idx, outdir=outdir, ID=ID, start=start)
# run the consistency test
consist_data_file, consist_result_file = MS.LiP_XL_MS_ConsistencyTest()
# select the representative structures from the consistency test
MS.select_rep_structs(consist_data_file, consist_result_file,
total_traj_num_frames=335, last_num_frames=67)
~~~ Outputs excel workbooks with summaries of the statistical tests for each meta-stable state and LiP-MS/XL-signal.
  b. Visualize representative structures from the ensemble.

#### Detect misfolding involving native entanglements using high-throughput experimental data and selecting candidates for further investigation

Timing (∼5 hrs. – days) depending on number of protein structures in ensemble and experimental signals. This section uses high throughput experimental data from a differential study which reports on the presence of conformational changes upon some kind of perturbation of the multi-protein sample. Here we use LiP-MS experiments done on proteome-wide samples that have been unfolded and refolded compared to untreated samples. This section is independent of the previous sections comparison to mass spectrometry data and therefore we renumber the protocol steps.

1. For all proteins that are observable in the high-throughput experimental data calculate the representative NCLE and their associated features using steps 1 – 8 in the “*Identify NCLEs in the native structure*” section of the protocol section.
2. Process the LiP-MS data to obtain a set of residues with statistically significant changes in conformation. (Box 3) *Critical Step*: There are many ways to process raw LiP-MS data to obtain signals of conformational changes and this processing can heavily influence any downstream statistical tests.
3. Define any confounding variables that need to be controlled for such as amino acid type or solvent accessibility.
4. For each protein generate a data frame where the number of rows is the number of residues in the proteins and the columns are the response variable (i.e. was the residue a site of a significant change in conformation -> 1 or not -> 0), the region of the protein in which the residue was found (entangled -> 1 or not ->0), and any confounding variables (in this case, amino acid composition and solvent accessibility).
5. Import, initialize, and use the ProteomeLogisticRegression class from EntDetect to model the odds of misfolding depending on what structural region of the protein it resides. This will require all the dataframe files made in step 4 of this section to be in a single directory, a file containing the list of UniprotID’s you want to analyze, and the regression formula written in the following format (y ∼ x_1_ + x_2_ + … + x_n_) following the stats model package style, (Box 4). ~~~
from EntDetect.statistics import ProteomeLogisticRegression
## Define some input paths and parameters
dataframe_files = “/path/to/dataframe/files”
outdir = “./population_modeling/”
gene_list = “/path/to/gene/list.txt”
ID = “Ecoli_noChaperones”
reg_formula = “cut_C_Rall ∼ AA + region”
## initialize the ProteomeLogisticRegression object
ProtRegession = ProteomeLogisticRegression(dataframe_files=dataframe_files, outdir=outdir,
gene_list=gene_list, ID=ID, reg_formula=reg_formula)
## Load the data into a dataframe for the regression
ProtRegession.load_data(sep=‘|’, reg_var=[‘AA’, ‘region’], response_var=‘cut_C_Rall’,
var2binarize=[‘cut_C_Rall’, ‘region’], mask_column=‘mapped_resid’)
## Run the regression analysis
reg_df = ProtRegession.run()
~~~
6. Take the exponential of the coefficient for a given term in the linear fit to obtain an odds ratio which can be used to judge the magnitude and sign of the association between the response and the regression variable when all other variables are held constant. The p-value of the coefficient quantifies how significant is this association. *Critical Step*. If this test is done on multiple sets of proteins (i.e. LiP-MS experiments done under different conditions, or on subsets of the sample) then you must apply an FDR correction to determine which associations are still significant after limiting the amount of false positives to less than 5%.
7. Import, initialize, and use the MonteCarlo class from EntDetect to select subpopulations of proteins with entanglements that are most likely to be misfolding prone. ~~~
from EntDetect.statistics import MonteCarlo
## Define some input paths and parameters
dataframe_files = “/path/to/dataframe/files”
outdir = “./monte_carlo/”
gene_list = “/path/to/gene/list.txt”
ID = “Ecoli_noChaperones”
reg_formula = “cut_C_Rall ∼ AA + region”
## initialize the clustering object
MC = MonteCarlo(dataframe_files=dataframe_files, outdir=outdir, gene_list=gene_list,
ID=ID, reg_formula=reg_formula)
## Load the data into the MonteCarlo object
MC.load_data(sep=‘|’, reg_var=[‘AA’, ‘region’], response_var=‘cut_C_Rall’,
var2binarize=[‘cut_C_Rall’, ‘region’], mask_column=‘mapped_resid’, ID_column=‘gene’,
Length_column=‘uniprot_length’)
## Run the Monte Carlo simulation
MC.run(encoded_df=MC.data, ID_column=‘gene’)
~~~ Outputs objective function and odds ratio statistics of each bin in the Monte Carlo scheme across the course of the simulation.
8. After the simulation has reached a steady state take the last 100 Monte Carlo steps and rank order the groups based on their average odds ratio. The group with the largest odds ratio contains proteins most likely to have experimentally observed changes in conformation that are strongly associated with their native entanglements.
9. To further refine the selection run multiple independent Monte Carlo simulations and select only those proteins that are in the largest odds ratio group more than 70% of the time.

### Anticipated outcomes

This protocol outlines an analysis workflow using a sample dataset (ecPGK) to identify NCLEs in experimental and computational structures (trajectories). It also showcases the examination of misfolded structural ensembles, analysis of folding pathways, extraction of representative structural ensembles consistent with LiP- and XL-MS data, and assessment of associations between entanglements and proteome-wide MS data. By following this protocol, users can produce the following key outputs:

1. A set of non-redundant NCLEs identified in the native ecPGK structure.
2. A set of non-redundant NCLE changes identified within the simulated, refolded ecPGK ensemble compared with the native structure.
3. Metastable states derived from the refolded ecPGK ensemble.
4. Lists of LiP-MS PK cut-sites and XL-MS crosslink residues exhibiting structural changes consistent with near-native metastable state ensembles obtained from simulation.
5. Representative structural ensembles of refolded ecPGK that display consistent structural changes as detected in LiP- and XL-MS experiments.
6. Odds ratios with corresponding p-values indicating associations between entanglements and misfolding observed in proteome-wide LiP-MS data.
7. A set of proteins likely to misfold due to their native entanglements, as suggested by proteome-wide LiP-MS data.

Due to the randomization features integrated into some steps of the protocol, users may not obtain identical outputs to those described here. For instance, the k-means clustering applied in MSM (see Procedure step 13) involves randomness, resulting in varying microstate assignments and consequently, slightly different metastable states. Nonetheless, the overall findings and conclusions remain robust and are not affected by this randomization.

### Trouble shooting table

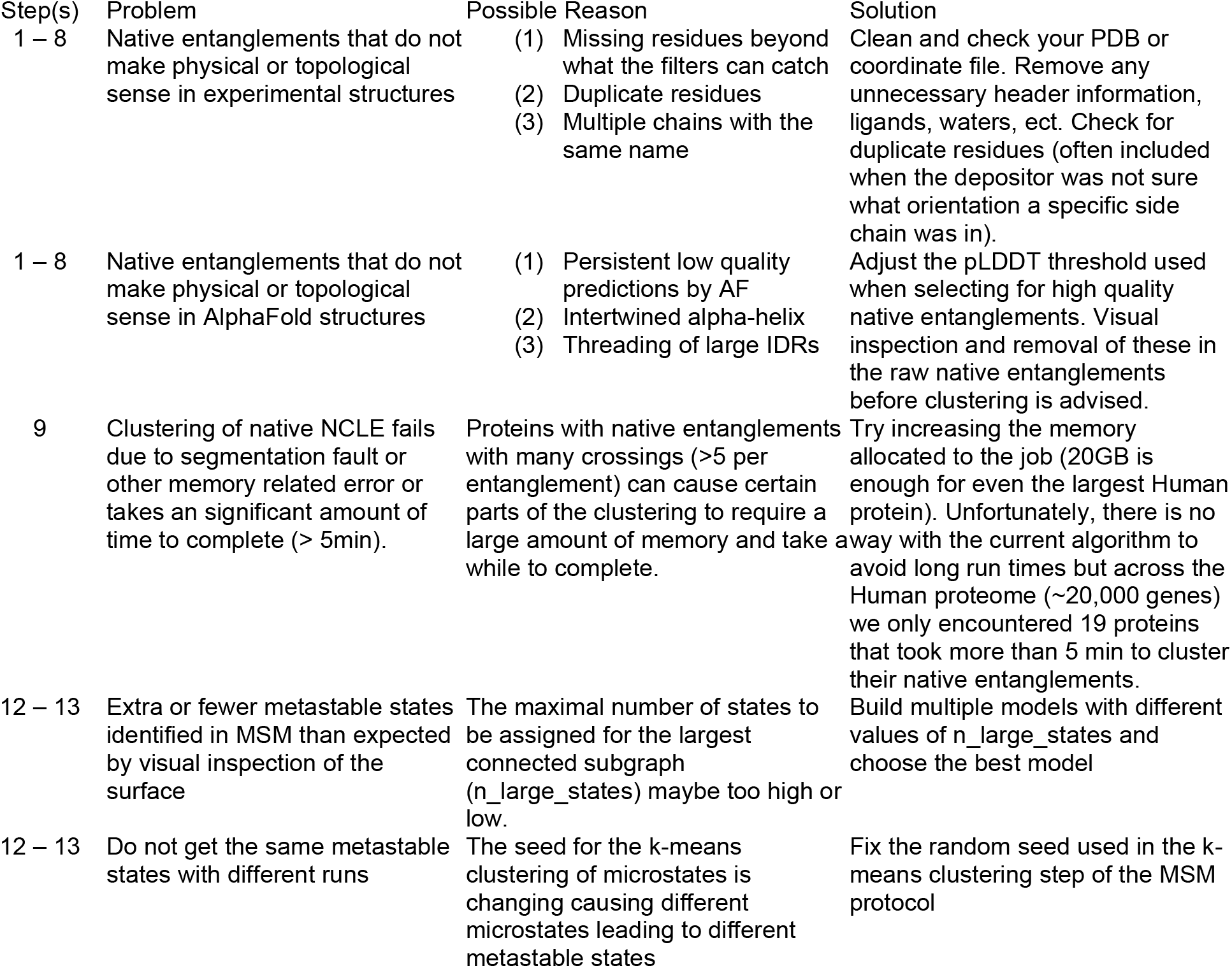
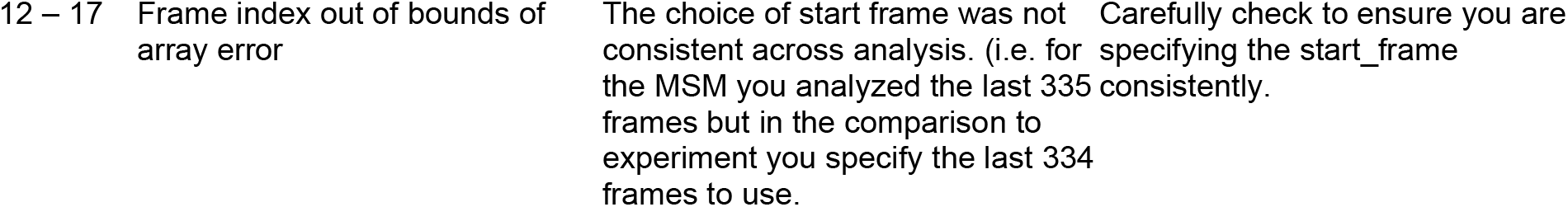

### Step Timings

- Identify NCLEs in the native structure (steps 1 – 8) ∼ Minimal time is less than 1 min but can take up to several hours depending on the size of the protein and number of raw entanglements to cluster.
- Identify changes in NCLEs and calculating other order parameters from simulation trajectories (steps 9 – 13) ∼ Depends on the depending on the size of the protein and length of trajectory. In our experience with medium sized proteins where the last 50 ns of each trajectory is analyzed it can take up to 2 days.
- Identify entangled structural ensembles consistent with high-throughput experimental data (steps 14 – 17) ∼ 1 to 2 day depending on the size of the protein and length of trajectory.
- Detect misfolding linked to native entanglements using high throughput mass spec and selecting candidates ∼ less than 1 min.
- Monte Carlo simulation for selecting candidates for further investigation ∼ 10 hrs to 2 days depending on the number of proteins and quality of experimental data.

### Box 1 Mathematics of identifying entanglements

For a given conformation of an *N*-residue long protein, a NCLE can be identified by first defining a set of loop forming native contacts as two residues (*i, j*) with alpha carbons (*C*_*α*_) within 8 Å of each other or heavy atoms within 4.5 Å of each other. Whether these loops are entangled with either or both terminus of the protein can be determined by examining the Gaussian Linking

Number (GLN) or the Topoly Linking Number (TLN). It is recommended to use TLN when possible as they are more accurate and have the added benefit of identifying the residues along each terminus that pierce the plane of a loop at the expense of computational speed. A combination approach is also beneficial where the GLN is first calculated to identify the subset of loop forming contacts that are potentially entangled and then the TLN is calculated for just this subset^5^.

#### Gaussian Linking Number (GLN)

Consider a loop closed by the residues (*i, j*) and the coordinates of the Cα atom in residue *l* (*r*_*l*_) of the protein, the partial linking values *g*_*N*_(*i, j*) and *g*_*C*_ (*i, j*) are then calculated using the discrete form of the Gaussian linking integral^65^ as:

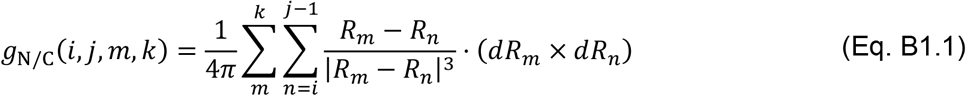

Where the midpoint between the two residues alpha carbons is 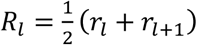, and the gradient is *dRa*_*l*_ = *r*_*l*+1_ − *r*_*l*_. Where *m* = 5 and *k* = *i* − 4 for the N terminus, and *m* = j + 4 and *k* = *N* − 6 for the C terminus. The loop is then considered to be entangled with either the N or C terminus if *g*_N/C_(*i, j, m, k*) ≥ 0.6. The GLN for the N and C terminus of the protein is then defined as the rounded *g*_N_ and *g*_C_ values where a non-standard rounding threshold^6^ of 0.6 is used instead of the more typical^3,4^ 0.5 (i.e. 0.58 -> 0, 0.63 -> 1, 1.58 -> 1, 1.63 -> 2). Finally to mitigate false positives or negatives caused by missing residues in a structure, entanglements with a loop where more than 5% of the residues are missing or more than 3 consecutive missing residues or where topoly fails to find a crossing residue (if topoly was used) should be discarded.

#### Topoly Linking Number (TLN)

Consider a loop closed by the residues (*i, j*), the residues along the N or the C terminus that pierce this loop can be derived from the Topoly^38^ python package which identifies the bond vectors which cross a tessellated loop spanning plane. This approach is much slower than the GLN method but more accurate in identifying loops that are truly entangled with their terminus. The TLN for a given terminus is then defined as the sum of crossing event chiralities^22^.

#### Entangled regions

For each representative NCLE (Box 2) three sets of key entangled residues can be defined:

i. The loop forming native contacts (*i, j*) and any crossing residues, *c*, identified by their TLN.
ii. The residues within +/-3 residues along the primary structure of the residues in set (i).
iii. The residues with (*C*_*α*_) within 8 Å of the residues in set (i) and (ii).

#### Change in NCLE status

The change of NCLE status refers to the change observed in the linking number (either GLN or TLN) for a loop in the target structure compared with a reference structure^3,4,22^. For example, the change from 0 to +1, or from +2 to -1.

#### Order parameter for changes in NCLE (G)

We can calculate *G* as the fraction of the native contacts where there was a change in either the N or C terminal entanglement status relative to a reference structure.

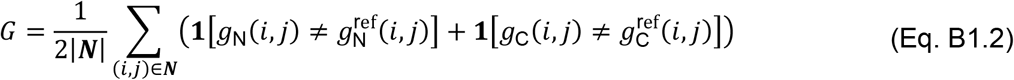

where ***N*** is an array containing all native contacts in the reference structure, (*i, j*) defines a specific loop closing native contact between residues *i* and *j* and **11**[·] is the indicator function which returns 1 when condition is satisfied and 0 otherwise. We initially proposed^3,4^ to consider changes in the total linking number, the sum of GLN or TLN for the N and C termini, which ignores the case where N and C termini have both changes in opposite direction (for example, N-terminus changes from 0 to 1 and C-terminus changes from 0 to -1, resulting a sum of 0, with no change) ^3,4,22^. We recommend considering the changes in each terminus separately^22^ as in Eq. B1.2.

### Controlling for slipknots and knots

To control slipknots the TLN can be used. If the TLN for a given terminus is 0 the entanglement with that terminus is disregarded as not a true pierced lasso. Proteins with knots are likely to also have a NCLE and it is useful sometimes to screen structures for their presence using either the Knotprot database^28^ or the Topoly package^38^.

### AlphaFold structures

AlphaFold (AF) structures^66,67^ with native entanglements have an additional set of criteria to reduce the inherent errors in the AF models from propagating to any downstream analysis.

1. The structure must have an overall <pLDDT> greater than or equal to 70.
2. The loop closing native contacts of an entanglement (*i, j*) must have a pLDDT >= 70.
3. The crossings must pass the following high-quality test:
  i. For a given termini that has an entanglement present start at the loop base (*i* or *j*) and in order examine each crossing.
  ii. If the first crossing has a pLDDT >= 70 it is kept and then you move onto the next.
  iii. The first instance of a crossing where the pLDDT < 70 you discard this crossing and any other crossings after it. (if the first crossing after the loop base is low-quality, we disregard the whole entanglement with that terminus)

### Box 2 Clustering native entanglements and their changes

#### Clustering native NCLEs in a single structure

For each protein structure there are many degenerate entanglements and therefore we seek to derive a set of unique representative entanglements through a clustering algorithm^6^ that has three main steps:

1. Cluster together loops that share the same crossings and chirality identified by topoly and choose the smallest loop to be the representative loop of each subcluster.
2. Then merge subclusters based on meeting three criteria.
  i. The crossings between the two sets of entanglements are spatially close.
  ii. Loops overlap to any extent.
  iii. No crossing from either entanglement considered is within the loops of another in the cluster.
3. Finally, subclusters are merged based on whether the distance between the geometric median of their representative native contacts and crossing residue indexes is less than a threshold defined by examining hand curated sets of entanglements.

The result is for each protein a set of unique entanglement(s) is identified.

#### Clustering changes in NCLEs in a pool of possible structures

The observed changes in entanglement status across a pool of structures are often degenerate. To identify a non-redundant set of unique entanglement changes, we developed a clustering algorithm^22^ consisting of the following steps:

#### Step 1: Encode Changes of Entanglement Status

Each change in entanglement status is encoded in the format L*C*, where L indicates the absolute value of TLN and C indicates chirality. Changes are represented as follows:

- Absolute TLN: no change (denoted as #), gain (+), or loss (-)
- Chirality: no change (#) or switch (∼)

For example, a loop gaining chirality-switched linkages from -1 to +2 at both termini is encoded as [L+C∼, L+C∼].

#### Step 2: Group by Change Type

Loops are first categorized based on identical encoded entanglement change status. Clustering is performed within each category to resolve structural degeneracies, as described in the following steps.

#### Step 3: Cluster by Crossing Residues (Agglomerative Clustering)

For both N- and C-terminal crossing residues, the pairwise distance 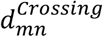 between loops *m* and *n* is computed as:

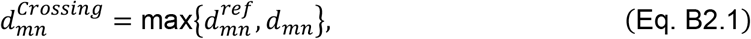

where 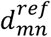 and *d*_*mn*_ are the absolute differences in median crossing residue indices between loops in the reference and simulation structures, respectively. If no crossing residues are present in a loop, the median is set to -1; mismatches between a loop with and without crossing residues are assigned a default distance of 10 residues. Clustering uses average linkage with a cutoff of 20 residues.

#### Step 4: Cluster by Loop Location (Agglomerative Clustering)

The loop location distance 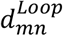 for loops *m* (*L*_*m*_) and *n* (*L*_*n*_) is calculated as:

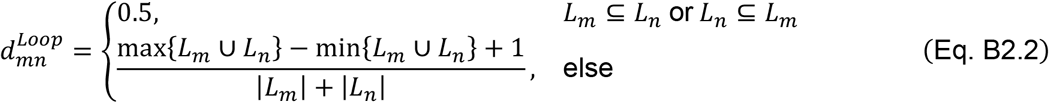

where |*L*_*m*_| is the length of the loop *m* and *L*_*m*_ ∪ *L*_*n*_ is the combined set of the loop residue indices. A distance of 1.0 corresponds to adjacent loops; values <1.0 indicate overlap, and >1.0 indicate separation. Clustering is done using average linkage with a cutoff of 1.0.

#### Step 5: Cluster by Crossing Contamination (Divisive and Agglomerative Clustering)

To separate two loops where one involves crossing residues of the other (*e*.*g*., loop (10, 20) is contaminated by the crossing residue 15 of another loop), the contamination distance 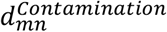 is defined as:

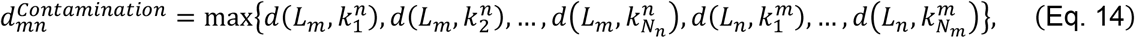

where 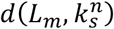 is the distance reflecting how deep the loop *L*_*m*_ formed by the native contacts (*i*_*m*_, *j*_*m*_) is contaminated by the *s*-th crossing residue 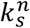 of the loop *L*_*n*_. It is defined as

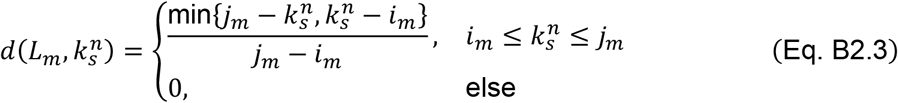

Divisive clustering is first applied to separate loops with contamination distances ≥ 0.1 (i.e., >10% loop length), followed by agglomerative clustering with average linkage and a cutoff of 0.1. We switched the linkage method from “complete” (as implemented in our previous work^22^) to “average” to better handle edge cases and produce more reasonable clusters. To minimize input-order bias, the *permuCLUSTER* algorithm^48^ is applied with 100 permutations in the abovementioned clustering steps 3, 4 and 5.

### Box 3. Preprocessing of LiP- and XL-MS Data

#### Processing Mass Spectrometry data at PK cut-site and crosslink site

To fit into our procedure, the Mass Spectrometry data must be preprocessed to remove degeneracy in the resulting signals used in downstream analysis (i.e. the PK cut-sites and cross-linked residues) and formatted to deliver essential information into a CSV file.

#### LiP-MS data

Ions are assigned and quantified with FragPipe^46^ and then processed with FLiPPR^68^, in which the peptides with the same PK cut-site are grouped together and the fold change of peptide intensities with FDR adjusted p-values (Welch’s t-test^69^ with BH correction^54^) are computed for each PK cut-site. Select PK cut-sites with adjusted p-value < 0.05 and put them in an Excel spread sheet. The final data file must have at least (1) “Cut Site” column including the cut-site residue of half tryptic peptides which are associated with PK cut-sites and (2) “Log2 FC” column including log2 fold changes of peptide intensities between refolded and native samples or any two conditions.

#### XL-MS data

The data processing involves a combination of label-free quantification using Proteome Discoverer^70^ and crosslink identification via XiSearch^51^/XiFDR^71^. Identified crosslinked peptides are mapped to features in the consensus file based on precursor m/z and retention time, allowing quantification across five replicates. For all replicates, missing values are imputed, and isotopically labeled pairs are matched by m/z offset and retention time. Peptides are filtered and merged at the crosslink residue level using the same criteria in FLiPPR^68^. The fold change of peptide intensities with FDR adjusted p-values (Welch’s t-test with BH correction) are computed for each crosslinking site. Select PK cut-sites with adjusted p-value < 0.05 and put them in an Excel spread sheet. The final data file must have at least (1) “Pairs” column including the crosslinked residues in a format of “K123-K456”, where “A” and “B” are amino acids followed by their residue numbers in the protein sequence and (2) “log2(heavy/light)” column including log2 fold changes of peptide intensities between refolded (heavy isotopically labeled peptide) and native samples (light isotopically labeled peptide). Details of the Mass Spectrometry data processing can be found in our previous publications^3,4,22,23^.

### Computing crosslinking propensity (XP)

The XP for a given residue pair is estimated using a modified version of the MNXL scoring function^22^, originally developed to evaluate crosslinking likelihood for Lys-Lys pairs based on solvent accessible surface distance (SASD)^55^. Since the original MNXL score is limited in handling non-lysine residues and may not account for variations in crosslinker length, we introduced a generalized formulation. The XP score for a residue pair (*j, k*) in structure *s* is defined as:

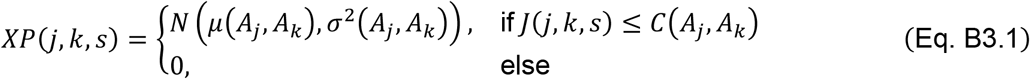

Here, *N*(*μ, σ*^2^) denotes the value of the normal distribution’s probability density function at distance *J*(*j, k, s*), with mean *μ*(*A*_*j*_, *A*_*k*_) and variance *σ*^2^(*A*_*j*_, *A*_*k*_), which are specific to the amino acid types *A*_*j*_ and *A*_*k*_. *J*(*j, k, s*) is the SASD between residues *j* and *k from structure s*, computed using the Jwalk algorithm^55^. *C*(*A*_*j*_, *A*_*k*_) is the amino-acid-specific cutoff distance for crosslinking. The mean and cutoff distances are adjusted to reflect differences in crosslinker length (specifically for DSBU in this protocol) and residue side chains. The standard deviation is scaled to maintain a fixed probability density at three standard deviations below the mean (*μ* − 3*σ*).

### Box 4 Modeling and ranking associations between misfolding and native entanglements in heterogeneous proteomics data

#### Controlling for differences protein abundance

Controlling for effects of endogenous protein abundance differences in the LiP-MS data can be done through the sum of the peptide ion abundances (SPA). The SPA for protein *i* is defined as:

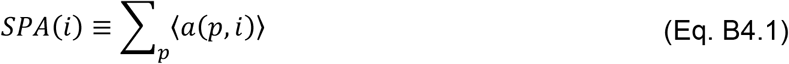

Where ⟨*a*(*p, i*)⟩ is the average peptide abundance reported for peptide *p* in protein *i* across the untreated technical replicates. The SPA cumulative density function can then be used to threshold the dataset based on the desired lower or upper bounds to test the robustness of results as a function of estimated protein abundance.

#### Logistic regressions

Logistic regressions^57,72^ are useful to model binary outcomes of a heterogenous proteomics dataset while controlling for confounding factors. At the protein level, the log-odds of a protein misfolding, defined as having at-least 1 half-tryptic peptide with a significant change in abundance between conditions, can be modeled as:

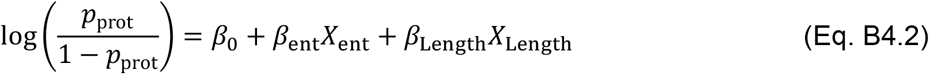

where *X*_ent_ is a binary variable defining whether the protein had a native entanglement and *X*_Length_ is the standard scaled length of the protein structure. An odds ratio (OR) for the odds of misfolding when a native entanglement is present relative to when it is not can then be estimated as 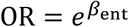.

The log-odds of observing a significant change in proteolysis susceptibility at a particular residue can be modeled as

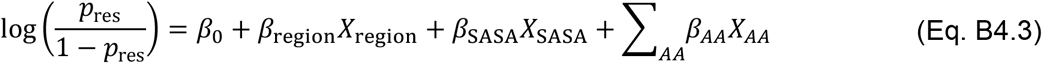

where *X*_region_ is a binary variable defining if the residue was in the entangled region of the protein or not, *X*_SASA_ is the standard scaled solvent accessible surface area of the residue, and *X*_*AA*_ is a binary variable defining if the residue was of the type *AA* where *AA* is one of the 20 canonical amino acids. An odds ratio for the odds of misfolding when a native entanglement is present relative to when it is not can then be estimated as 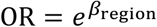.

#### Selecting populations of proteins likely to misfold involving their native entanglement

Given a set of heterogenous set of proteins observed in a differential proteomics experiment where the statistical power is too weak to discern changes in some feature of a single protein we can at the very least rank or subgroups of proteins that are most likely to exhibit the feature change relative to the other groups. For example, a set of proteins can be randomly distributed amongst 4 groups of equal size. For each protein group *i*, there is an objective function defined as:

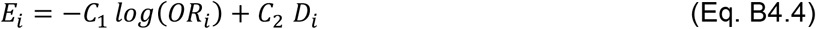

where *OR*_*i*_ is the ratio of the odds of misfolding in the entangled region relative to the odds of misfolding in the non-entangled region of the protein group *i* and *D*_*i*_ is the Kolmogorov–Smirnov test statistic^73^ comparing the protein size of group *i* to a reference distribution taken as the superset of all four groups, and *C*_1_ & *C*_2_ are arbitrary constants. One MC step is taken by randomly swapping 5 proteins between each pair of neighboring group indexes.

The Metropolis criteria^74^ is applied to determine if any given swap is accepted or rejected. The acceptance ratio for a swap between groups *j* and *k* at MC step *x* is defined as:

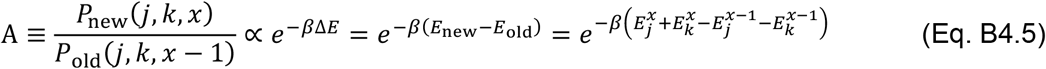

The swap is accepted if either A ≥ 1 or 1 > A > *u*, where *u* is a random number sampled from a uniform distribution bounded by [0,1]. β is an inverse temperature scaling factor that starts very low at 0.05 to allow ample room for the simulation to explore the objective function landscape and is scaled linearly to 1000 after every 750 MC steps and remains constant at 1000 thereafter.

## Acknowledgements

E.O. gratefully acknowledges support from the National Science Foundation (MCB-2031584) as well as from the National Institutes of Health (R35-GM124818). Portions of numerical computations and data analysis in this work have been carried out on high-performance computing collectively known as ROAR, which is operated by the Institute for Computational and Data Sciences at The Pennsylvania State University. The National Science Foundation (DBI-2335029) National Synthesis Center for the Emergence of Molecular and Cellular Sciences.

## Credit authorship contribution statement

**I.S**.: Methodology development, Formal analysis, Visualization, Validation, Data curation, Writing – original draft, Writing – review and editing.

**Y. J**.: Methodology development, Formal analysis, Visualization, Validation, Data curation, Writing – original draft, Writing – review and editing.

**E. O**.: Conceptualization, Formal analysis, Funding acquisition, Investigation, Methodology development, Project administration, Resources, Supervision, Validation, Writing – original draft, Writing – review and editing.

