## Supplemental Information 1 for "Detecting misfolded non-covalent lasso entanglements in protein structures, simulation trajectories, and mass spectrometry data"

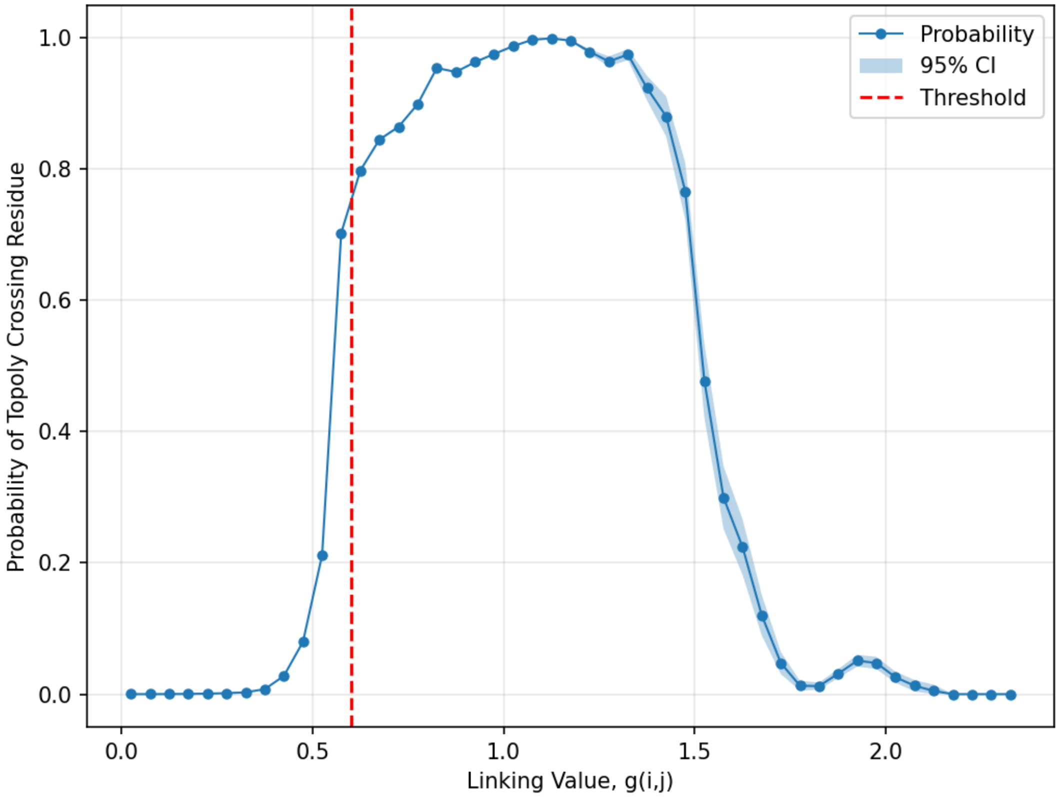


**Supplementary Figure 1:** The probability of the Topoly python package finding a crossing residue as a function of the absolute value of the Gauss linking value across the last 10 ns of all 1000 trajectories of ePGK. The Topoly default value for the min distance between two crossings was used (10 residues), but we use our terminal and loop buffers of 5 and 4 residues respectfully. Shaded regions are 95% confidence intervals. Red dashed line is the 0.6 threshold for determining the presence of a NCLE. At high values of $|g(i,j)|$ the default min distance between crossings of 10 residues dramatically decreases the accuracy. This parameter is variable in our EntDetect package and we suggest either optimization or using 5 residues instead.
